# Modeling behavior to disentangle motion-related effects in functional ultrasound imaging in awake, head-fixed mice

**DOI:** 10.64898/2026.02.04.703701

**Authors:** C. Qin, F. Nelissen, R. Waasdorp, A. Lotfi, C. Rojas, L. De Angelis, M. Heemskerk, D. Maresca, C. Keysers, B. Heiles, V. Gazzola

## Abstract

Functional ultrasound imaging (fUSI) enables brain-wide mapping of hemodynamic activity in awake rodents, offering a powerful bridge between systems neuroscience in animals and human neuroimaging. However, extending fUSI beyond tightly controlled sensory paradigms to more naturalistic behaviors is limited by motion-related confounds arising from locomotion, physiological arousal, and movement-linked neural activity. Here, we introduce a behavior-informed modeling framework that explicitly incorporates continuous measurements of running speed and head motion into a general linear model to account for motion-related variance while preserving interpretable task-related signals. We validate this approach in two head-fixed paradigms with distinct motion profiles: a visual stimulation task with minimal stimulus-driven movement, and a noxious stimulation task in which locomotion and arousal are intrinsic to the behavioral response. In both cases, explicit behavioral modeling recovers neural response patterns that more closely resemble low-motion reference conditions than blind, model-free denoising approaches. Critically, during noxious stimulation, behavior-informed modeling preserves shock-intensity-dependent activity in the primary somatosensory cortex that is attenuated by global component removal. These findings demonstrate that explicit modeling of behavior enables reliable interpretation of brain-wide fUSI signals during naturalistic, high-motion conditions, opening the door to studying affective and cognitive processes that were previously difficult to access with functional ultrasound imaging.

## Introduction

fMRI has become the dominant tool to study large-scale organization in the human brain, which infers neural activity from blood-oxygenation changes across the whole brain (Ogawa et al., 1992). Meaningful integration of pre-clinical and human studies requires direct comparisons of brain activity to evaluate the translational validity of animal models. While small-bore animal MRI scanners offer powerful insights into brain function, their confined space and loud acoustic environment make them unsuitable for studying complex and naturalistic behaviors. This poses a major limitation for rodent research in cognitive and affective neuroscience, where experiments often involve head-fixed mice performing tasks like navigation, decision-making, or social interaction, sometimes in virtual reality settings (Chari et al., 2023; Krishnan et al., 2025; Stuart et al., 2024).

Functional ultrasound imaging (fUSI) holds the promise to overcome some of these limitations and thereby enable more principled cross-species comparisons. fUSI detects blood volume variations in the cerebral vascular space, and, much like BOLD fMRI, can infer brain activity through the neurovascular coupling (Macé et al., 2011). Physically, fUSI measures the intensity of the Doppler signal, which is proportional to the cerebral blood volume and is independent of the vessel distribution, orientation, and flow velocity (Montaldo et al., 2022). Because ultrasound penetration depends on the frequency of the waves, acoustic waves of different frequencies can be used to image a wide range of animals from zebrafish (Chang et al., 2019), rodents (Macé et al., 2011; Tiran et al., 2017) and ferrets (Bimbard et al., 2018) to non-human primates (Dizeux et al., 2019; Norman et al., 2021; Takahashi et al., 2021) and humans (Imbault et al., 2017; Rabut et al., 2024; Soloukey et al., 2020). Its attractive spatiotemporal resolution (100ms, 100um at 15MHz), and its relative portability allow to image brain activity in freely moving (Tiran et al., 2017), head-fixed (Edelman et al., 2024) and sleeping models (Bergel et al., 2024, 2025), making it suitable for studying brain-wide dynamics during complex rodent behavior. Furthermore, fUSI is compatible with other modalities such as electrophysiology (Nunez-Elizalde et al., 2022; Zhang et al., 2024) and optogenetics (Aurup et al., 2023; Brunner et al., 2020; Edelman et al., 2021; Provansal et al., 2021; Rungta et al., 2017; Sans-Dublanc et al., 2021), allowing preclinical measurements across multiple spatiotemporal scales.

Extending fUSI beyond exogenous-stimulus paradigms into domains such as decision making and affective neuroscience is a critical step toward understanding how neural activity gives rise to internal states and behavior. In contrast to classical stimulus-locked paradigms that primarily target sensory processing, these domains emphasize interactions between external stimuli and internal cognitive and emotional states, shaping context-dependent behavioral responses through distributed, brain-wide networks. This shift, however, requires addressing a fundamental challenge: the pervasive influence of motion-related confounds. Because fUSI signals are derived from the backscatter of ultrasound waves by red blood cells, even minor displacements of the brain can cause substantial artifacts in the reconstructed images. Unlike optical preparations with miniaturized, implanted two-photon microscopes (2P; Zong et al., 2017), current fUSI probes are typically external in head-fixed mice, so small brain–probe displacements are more consequential. Standard in-plane registration (as used in 2P) (Pnevmatikakis & Giovannucci, 2017) can improve fUSI data, but a substantial fraction of fUSI motion is out-of-plane and coupled to the acoustic field, which cannot be fully corrected by conventional image-based methods. In addition to the direct mechanical artifact during locomotion, whether head-fixed on a treadmill or freely moving, systemic hemodynamic changes such as increased cardiac output and arterial pulsatility, could modulate cerebral blood volume independently of neural activity, particularly in large arteries, thereby confounding the interpretation of localized neural activity. Finally, overt movements themselves engage motor and premotor circuits, generating genuine neural signals that correlate with motion. Together, these three facets of the motion problem - mechanical distortion, systemic physiology, and neural co-activation - make it challenging to isolate stimulus- or cognition-related fUSI signals in behaving animals.

Several approaches have been developed to mitigate motion-related confounds in fUSI data. Spatiotemporal clutter filtering (Demene et al., 2015; Urban et al., 2015) removes signals that are either too static to reflect blood flow, such as from the skull, or fluctuate too rapidly to be physiologically meaningful, thereby isolating relevant vascular activity in each frame, or power Doppler image (PDI). Subsequent steps often involve discarding and interpolating frames contaminated by large movement (Brunner et al., 2021) or excluding entire epochs (Brunner et al., 2020). In addition, removing the first principal component (PC) from the spatiotemporal decomposition of the entire time series has been used to further suppress motion-related artifacts (Edelman et al., 2021), and more recently, data-driven decomposition approaches have been extended to further separate motion-related components from neural signals in fUSI time series(Meur-Diebolt et al., n.d.-a). While these methods improve signal stability, they are largely signal-driven and uninformed by the animal’s actual movements. As a result, they suppress variance based on statistical or spatiotemporal properties rather than explicitly modeling motion as a measurable variable. Consequently, potentially informative neural–behavior relationships may be attenuated or removed, and residual motion-related artifacts can persist in dynamic behavioral paradigms.

Building on these existing methods, we present a complementary modeling framework that explicitly incorporates continuous recordings of running speed and head motion to account for movement-related variance in fUSI data. By integrating these behavioral covariates into a general linear model (GLM), motion-linked components are included as regressors of no interest - analogous to the inclusion of motion parameters in task-based fMRI GLMs - while preserving interpretable neural variance associated with experimental conditions. This framework enables us to quantify the contribution of movement-related signals to the fUSI time series and to recover stimulus-evoked or affective responses more faithfully in awake, behaving mice.

We validated this framework across two head-fixed experimental paradigms: a well-characterized visual stimulation paradigm which should not elicit stimulus-driven body motions, but only present spontaneous (loco-)motion, and a noxious stimulation paradigm that induces substantial locomotion and physiological arousal as a consequence of the stimulus. By comparing performance across these conditions, we demonstrate that motion-induced variance can be effectively modeled and separated without discarding large portions of data, outperforming a PCA-based approach. Incorporating behavioral regressors substantially reduces motion-related confounds while preserving biologically meaningful task-response patterns. Together, these findings show that fUSI, when combined with explicit behavioral modeling, can support more naturalistic experimental designs in head-fixed mice, bridging the gap between controlled stimulus paradigms and the study of complex, self-generated behavior.

## Results

### Characterization and correction of motion in fUSI data

To assess how motion affects fUSI signals in awake mice, we first tested our method in a visual stimulation paradigm, which is commonly used in neuroscience (Gesnik et al., 2017; Macé et al., 2018; Maresca et al., 2020), as a proof of concept. We then applied a noxious stimulation paradigm, which complicates the separation of motion- and stimulus-related activity, to test our ability to dissociate overlapping signals. Awake, head-fixed mice were placed on a running wheel (Fig. 1A), while fUSI data were acquired with a linear probe through a cranial window (Fig. 1AB). A recording session consisted of a short baseline period followed by 5-10 visual stimulation blocks (15 s on, 45 s off) with gratings at eight random orientations (Fig. 1C; Table S1). In the final three sessions for each animal, tail shocks were subsequently delivered in 8 blocks of three 2 s shocks, with intensity per block progressively increasing from 0.05 to 0.4 mA in 0.05 mA steps, with five 0 mA control trials before, midway, and after the session (Fig. 1D; Table S1). To characterize and mitigate motion-related artifacts in the fUSI data, the animal’s body movements were quantified: running speed was continuously monitored using a wheel-mounted encoder on the treadmill and head motion was concurrently recorded with a three-axis accelerometer affixed to the head plate (Fig. 1A). In this section, we first characterize the influence of motion on the high-pass–filtered fUSI data in both experiments, and then examine how motion-related artifacts can be mitigated using different analytical approaches.

**Figure 1.**
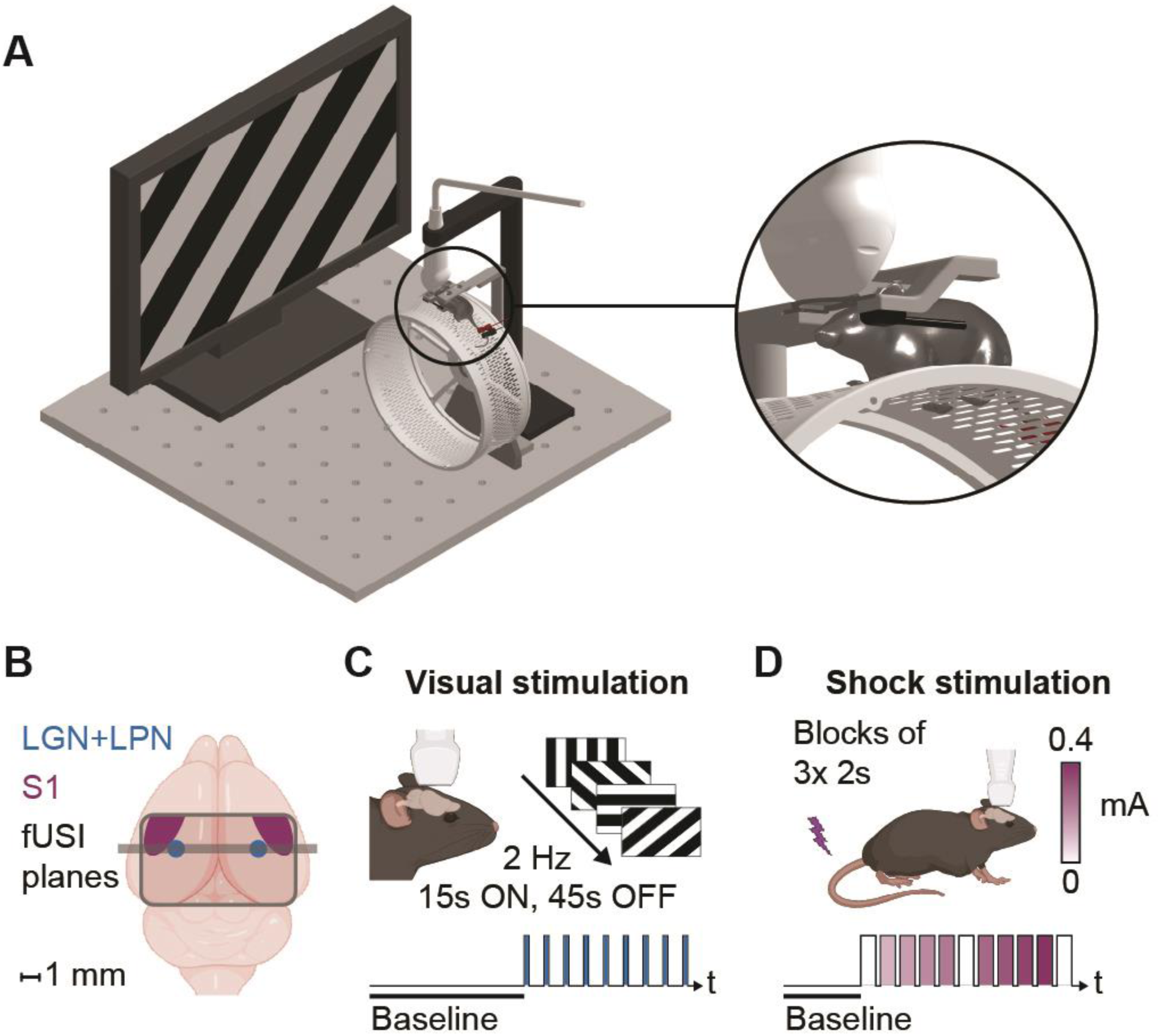
Experimental paradigm, characterization, and accounting for motion-related effects. (A) Experimental configuration with monitor for visual stimuli presentation and tail shock electrodes for shock administration. Motion monitoring is performed via a running wheel (grey) and an accelerometer underneath the headplate holder (black) along with fUSI recordings acquired using a Verasonics L22-14vX High Frequency Imaging probe (central frequency 18.5 MHz; white) (B) Cranial window and fUSI plane containing the lateral geniculate nucleus and lateral posterior nucleus of the thalamus (LGN + LPN, blue) and primary somatosensory cortex (S1). (C) Experimental timeline of one visual stimulation session: After a 2-5 min baseline (black screen), 5-10 moving gratings (black/white, 8 angles, 2 Hz) were shown for 15 s, with 45 s inter-stimulus intervals (black screen) on a monitor ∼20 cm in front of the animal. Blue bars represent one 15 s stimulus period. 27 sessions across 5 animals were recorded (See Table S1). (D) Experimental timeline of one shock stimulation session: Mice received three tail shocks per intensity, increasing from 0.05 to 0.4 mA in 0.05 mA steps with 40 ± 10 s inter-trial intervals. Blocks of five control trials (0.0 mA) were delivered before the first shock, after the 0.2 mA shocks, and after the final shock. Purple bars represent three shocks of one intensity. 12 sessions across 5 animals were recorded (See Table S1).

### Characterization of motion in fUSI data

Mice exhibited distinct running behavior across the visual and noxious stimulation experiments. While the proportion of time spent running (speed > 2cm/s) did not differ significantly between the two experiments (visual: 0.40 ± 0.23, noxious: 0.30 ± 0.10, mean ± std, Fig. S1A), the average running speed during shock stimulation sessions was higher than during visual stimulation (visual: 16.96 ± 4.08 cm/s, noxious: 28.83 ± 6.09 cm/s, Fig S1B), reflecting a locomotor response to aversive stimuli. We then estimated the amount of variance explained by the different motion parameters in the fUSI data (Fig. 2D) using a General Linear Model (GLM). While head motion explained only marginal variance in the visual experiment, with R^2^ = 0.0064 ± 0.0075 its contribution was substantially larger in the noxious experiment: R^2^ = 0.1073 ± 0.0632. Running speed irrespective of whether it was convolved with the hemodynamic response function or not, contributed to a significant amount of variance in the fUSI signal, jointly accounting for 0.11 ± 0.048 and 0.25 ± 0.048 of the total variance in each experiment. The variance explained by the acceleration (Running’, i.e. first derivative or running speed) was negligible (0.00016 ± 0.00019 in the visual stimulation, 0.0053 ± 0.0024 in the noxious stimulation), and was therefore omitted from subsequent analyses.

**Figure 2.**
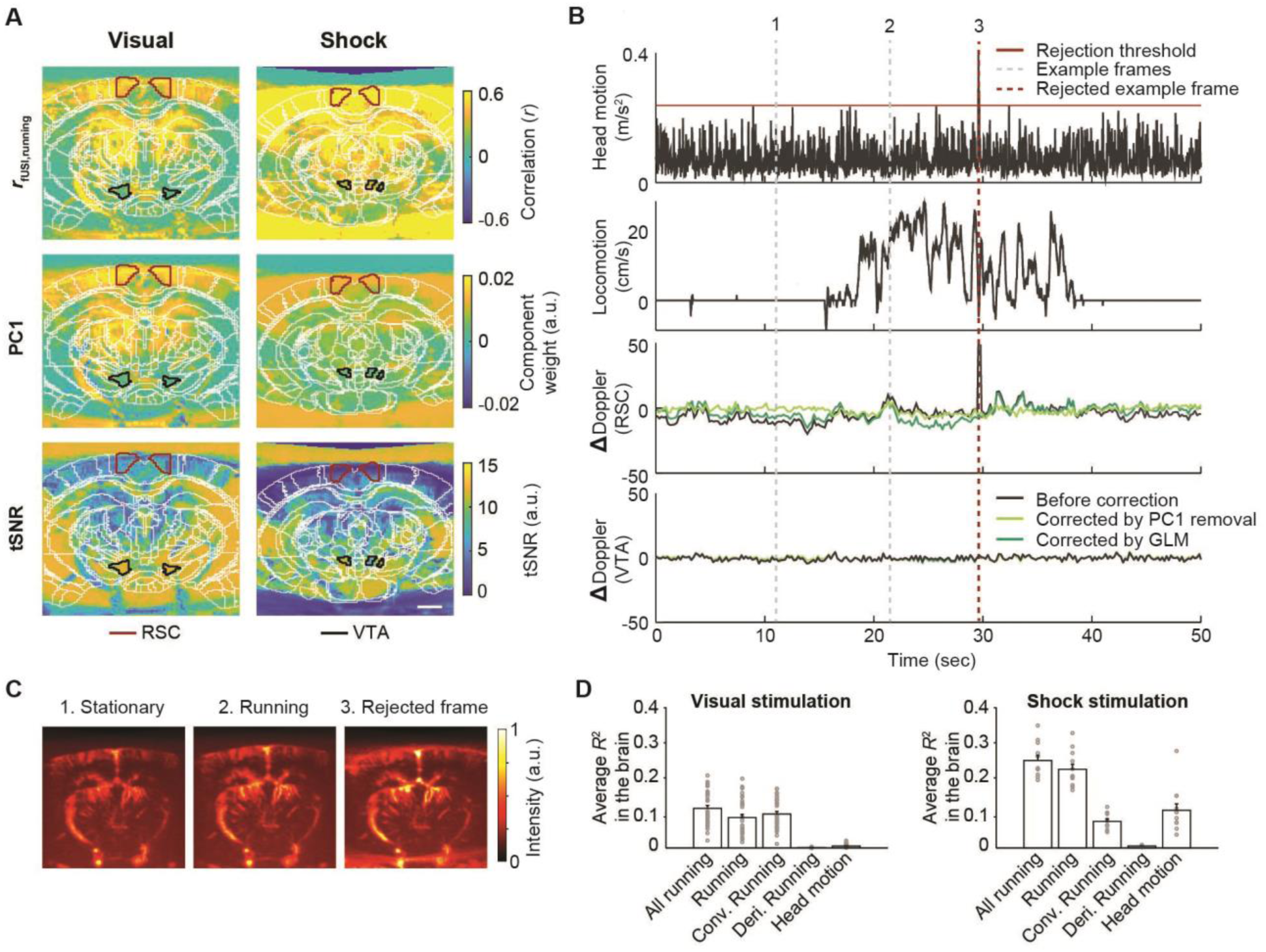
Characterization and accounting for motion-related effects. (A) Spatial maps of the correlation between running and fUSI signal, weight of principle component 1 (PC1), and temporal signal-to-noise ratio (tSNE) in each pixel during the baseline, visual-stimulation period and noxious-stimulation period of a representative example session. (B) Head-plate acceleration, running wheel velocity, and the Power Doppler signal from selected regions before (black) and after handling motion-related effects using PC1 removal (light green) or GLM-based modeling (dark green) methods. RSC = retrosplenial cortex, VTA = ventral tegmental area. (C) Example Power Doppler Images (PDIs) corresponding to time points 1, 2 and 3 shown in panel F with dotted lines, showing instances when the animal was stationary and running, and one rejected frame. (D) Variance explained by different motion variables in visual stimulation and noxious stimulation paradigms.

Given that running contributes substantially to widespread variance in the fUSI signal (Fig. 2AD), we next examined how this behaviorally related variance overlaps with the dominant shared component captured by principal component analysis (PCA) in both experiments. The first principal component (PC1) captures the largest portion of variance shared across many pixels, and is often interpreted as reflecting motion-related artifacts and other global fluctuations, but can also include genuine neural signals that are widespread or shared across networks (Gotts et al., 2020). Here, PC1 explained 0.51 ± 0.082 and 0.73 ± 0.13 of the total variance in the visual and noxious stimulation experiments, respectively, and the spatial pattern of PC1 was similar to the running correlation map (visual: r = 0.52 ± 0.32, shock: r = 0.90 ± 0.11), indicating that the variance explained by the running variables are also partially captured by the PC1 (Fig. 2A).

Additionally, to quantify temporal signal stability in the fUSI signal, the temporal signal-to-noise ratio (tSNR) was computed for each pixel (Fig. 2A). tSNR was anticorrelated with the fUSI-running correlation map (visual: r = -0.45 ± 0.14, shock: r = -0.67 ± 0.08), indicating reduced signal reliability in regions more strongly correlated with running.

### Accounting for motion-related effects in fUSI data

Following this characterization, we next address the mitigation of motion-related effects using a series of analytical procedures. First, fUSI frames with abrupt high-amplitude artifacts, caused by sudden movements, were detected by identifying frames with either substantial head movements measured by the accelerometer (session-wise *z* > 3.5; Fig. 2B panel 1) or extreme deviations in the fUSI signal at individual pixels (*z* > 5). Frames meeting either criterion were removed and interpolated (Fig. 2C; see method section ‘Preprocessing of fUSI data’). On average, this procedure resulted in a rejection rate of 0.024 ± 0.0014 of all frames in the visual-stimulation experiment. Consistent with the greater variance explained by head motion in the noxious condition (Fig. 2D), the rejection rate in this experiment was slightly higher, at 0.028 ± 0.0011 (*t*_(37)_ = 7.52, *p* = 5.83*10^-9^).

We subsequently addressed motion-induced signal changes using two alternative approaches: by removing PC1 or by including the running speed measurements in a GLM and examining the residual signal (Fig.1D). To illustrate the effect of these different approaches to handling motion-related effects, we selected two regions of interest (ROIs): the restrosplenial cortex (RSC; Fig. 2B, panel 3), a region involved in locomotion (Mao et al., 2020), and the ventral tegmental area (VTA, Fig 2B, panel 4), which is not expected to be modulated by motion. The VTA thus serves as a negative control to ensure that motion-handling procedures do not introduce spurious activity. Removing the running speed via a GLM and removing PC1 altered the temporal structure of the signal in different ways. Throughout the following sections, we systematically evaluate analyses with and without PC1 removal, allowing us to assess how PCA-based denoising interacts with GLM-based modeling of motion-related effects in reducing artifacts while preserving neural signals.

### GLM modeling isolates visual responses from running-related variance

To assess the effectiveness of the GLM approach in disentangling visual-evoked activity from running-related signals in the visual stimulation experiment, we compared three models (Fig. 3C). First, we created a reference map by estimating visual responses during trials when the animals were stationary (M1). Consistent with their role in visual processing (Brunner et al., 2020), the lateral geniculate nucleus (LGN) and the lateral posterior nucleus of the thalamus (LPN) showed strong activation with this predictor (Fig. 3C, Visual(Stat), M1). Second, we modeled all visual stimulation trials (M2), including those in which the animals were running. As a result, the measured responses reflected a mixture of visual-evoked activity with running-related signals, introducing additional variability into the maps. In a representative session, this resulted in higher effect sizes across broader regions beyond the visual areas (Fig. 3C, Visual(All), M2). A similar pattern was observed using a correlation-based analysis that included all visual stimulation trials, with responses again extending beyond visual areas (Fig. 3C, Visual(All), Corr). In contrast, the third model (M3) included both running and a Visual × Running interaction term, capturing state-dependent modulation of visual responses by running and allowing estimation of visual-stimulus-related activity (Troncoso et al., 2015). This model showed high effect sizes for the visual predictor that were spatially localized to the LGN and LPN, with minimal activity in unrelated regions (Fig. 3C, Visual(All), M3).

**Figure 3.**
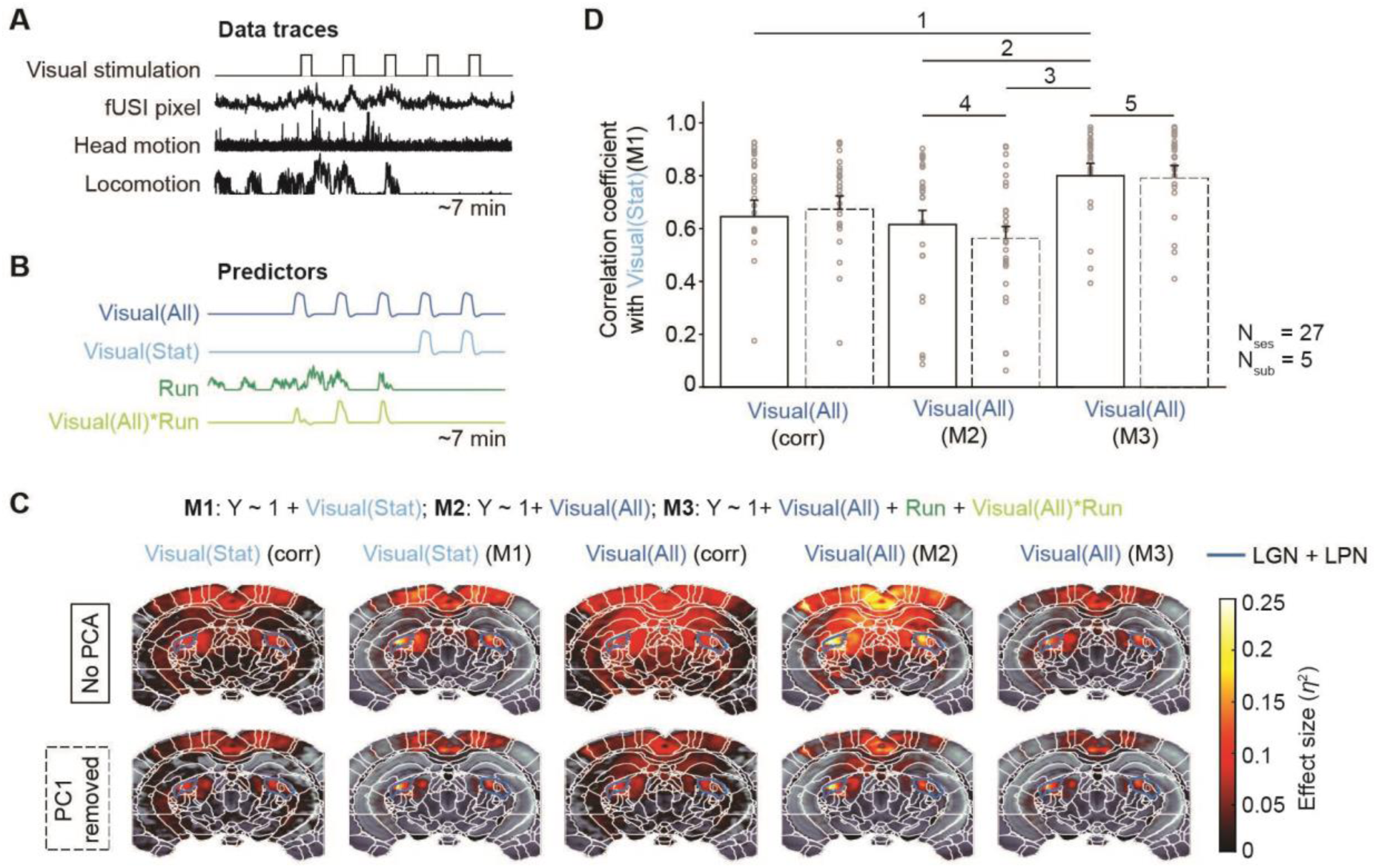
GLM modeling reveals visual responses independent of running. (A) Data traces of an example session depicting five visual stimuli, the signal from one fUSI pixel from the lateral geniculate nucleus (LGN), head-plate acceleration and running wheel velocity. (B) Example predictors used in the GLM, representing all trials (Visual(All), dark blue), stationary trials (Visual(Stat), light blue), Running (Run, dark green) and the interaction between running and the visual stimuli (Visual(All)*Run, light green). Run refers to Running and Running⊗HRF combined. (C) GLM definitions using predictors from B, and example effect size maps of predictors across different GLMs and correlations without PC1 removal (top row) and with PC1 removal (bottom row). Lateral geniculate nucleus and lateral posterior nucleus of the thalamus (LGN, LPN) are outlined in dark blue. (D) Group-level 2D correlations of effect size maps across GLMs and relative to the stationary visual trials, considered here the ground-truth model, shown without PC1 removal (solid lines) and with PC1 removal (dashed lines). Error bars represent standard deviation across sessions. Statistics represented in Table 1.

**Table 1.** Statistics of visual stimulation experiment. The *t*-values are from two-sided paired t-tests corresponding to Figure 3D. *BF_10_* indicates Bayes factors in favor of the alternative hypothesis. n = 27 for all comparisons.

| Comparison | Statistic |
| --- | --- |
| D1 | $t(26) = 6.45, p < 0.001, BF_{10} = 2.3 * 10^4$ |
| D2 | $t(26) = 6.87, p < 0.001, BF_{10} = 5.9 * 10^4$ |
| D3 | $t(26) = 7.86, p < 0.001, BF_{10} = 5.5 * 10^5$ |
| D4 | $t(26) = -2.16, p = 0.04, BF_{10} = 1.48$ |
| D5 | $t(26) = -0.66, p = 0.51, BF_{10} = 0.25$ |

To assess how well this method separated visual from motor responses on a group level, we treated the effect size map from stationary trials (Fig. 3C, Visual(Stat), M1) as a reference estimate of visual activity minimally influenced by running. We used a two-dimensional correlation to test whether including running in the full model (M3) produced visual effect size maps that were more similar to the stationary-trial reference (M1) than those obtained with the model without running predictor (M2). Formally, we tested whether:

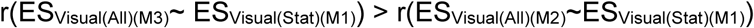

Additionally, we compared the visual effect size maps from M3 to those obtained using a simple correlation-based approach, by asking whether:

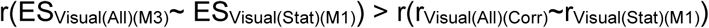

In line with these comparisons, the effect size map of the visual predictor from M3 showed the highest correlation with the stationary-trial reference (*r* = 0.81 ± 0.21), indicating the closest correspondence to the stationary-trial reference estimate of visual activity (Fig. 3D). In comparison, M2 had a lower correlation (*r* = 0.61 ± 0.28), while the correlation-based method using all visual trials showed an intermediate correlation that was significantly lower than that of M3 (*r* = 0.67 ± 0.27; Fig. 3D). Together, these results indicate that explicitly modeling running and its interaction with visual stimulation in the GLM (M3) yields visual response estimates that more closely resemble those obtained under stationary conditions, providing a more accurate estimation of neural responses to simple sensory stimuli in awake, behaving animals.

Because variance related to running is partially captured by the first principal component (PC1; Fig. 2A), we next examined how removing PC1 during preprocessing (see Methods) affects GLM-derived effect size maps. To this end, we repeated the GLM analyses after PC1 removal, allowing a direct comparison between blind, model-free denoising and behavior-informed modeling. The example effect size maps of stationary visual trials (Fig. 3C, Visual(Stat), M1) were highly similar with and without PC1 removal (*r* = 0.86 ± 0.16), demonstrating that removing PC1 did not substantially alter the effect-size maps related to visual stimuli, apart from a modest reduction in the spatial extent of some effect-size regions (Fig. 3C, lower row).

Next, to quantitatively evaluate whether removing PC1 during preprocessing improves visual response estimation in the presence of running, we compared the correlation of M2 effect-size maps with the stationary-trial reference (M1) before and after PC1 removal. Removing PC1 did not improve the correlation with the stationary-trial reference (M1); instead, correlations were slightly lower after PC1 removal than before it (Fig. 3D). This suggests that, while PC1 removal can reduce shared variance, it does not necessarily enhance estimation of visual responses when body movement is involved. Similarly, for the model that explicitly accounted for running (M3), removing PC1 did not improve visual response estimation: correlations with the stationary-trial reference were comparable with and without PC1 removal for Visual(All) of M3 (Fig. 3D). This indicates that explicitly modeling running renders the estimation of visual responses largely insensitive to this PC1 removal step.

Finally, we asked whether explicit behavioral modeling is more effective than blind source separation (PC1 removal) during preprocessing for separating visual responses from motion-related signals. To this end, we compared:

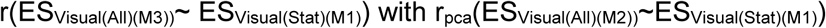

We found that explicit behavioral modeling yielded visual effect size maps more similar to the stationary-trial reference than those obtained using blind source separation alone, indicating superior recovery of visual response patterns (Fig. 3D).

While the analyses described above focused on spatial specificity, we next asked how different approaches to accounting for motion-related variance affect the detectability of stimulus-evoked signals by quantifying the contrast-to-noise ratio (CNR). The CNR was calculated in target regions (LGN and LPN) and a control region (VTA) before and after GLM-based modeling, after PC1 removal, and after combining both approaches (Fig. S2). Both the GLM and PC1 removal caused a slight decrease in the CNR in the target regions compared to the data prior to accounting for motion-related effects. The CNR did not differ between the two approaches, while combining both approaches resulted in a further reduction in CNR compared to each method alone. This effect was less pronounced in the control region: in the VTA, the CNR remained unchanged after the GLM, but decreased after PC1 removal. This indicates that the GLM approach and PC1 removal achieved a similar degree of preserving stimulus-evoked contrast in target regions, while PC1 removal additionally reduces variance in non-target regions, reflecting its global, model-free denoising of shared variance.

### GLM modeling dissociates shock-related neural responses from running-related variance

To test our ability to disentangle overlapping signals, we designed a challenging scenario using a noxious stimulation paradigm (Fig. 1D). Because tail shock stimulation reliably triggers escape and running behaviors (Krishnan et al., 2025), it is critical to dissociate neural activity associated with directly shock-related processing from activity related to these nocifensive movements. As expected, tail shocks reliably evoked robust locomotion time-locked to shock delivery (Fig. 4A), scaling with intensity (Fig; 4B) and generating pronounced motion artifacts (Fig. 2AD). Because locomotion temporally overlapped with stimulus delivery, we assessed how effectively the GLM approach reduced the impact of running on the measured responses.

**Figure 4.**
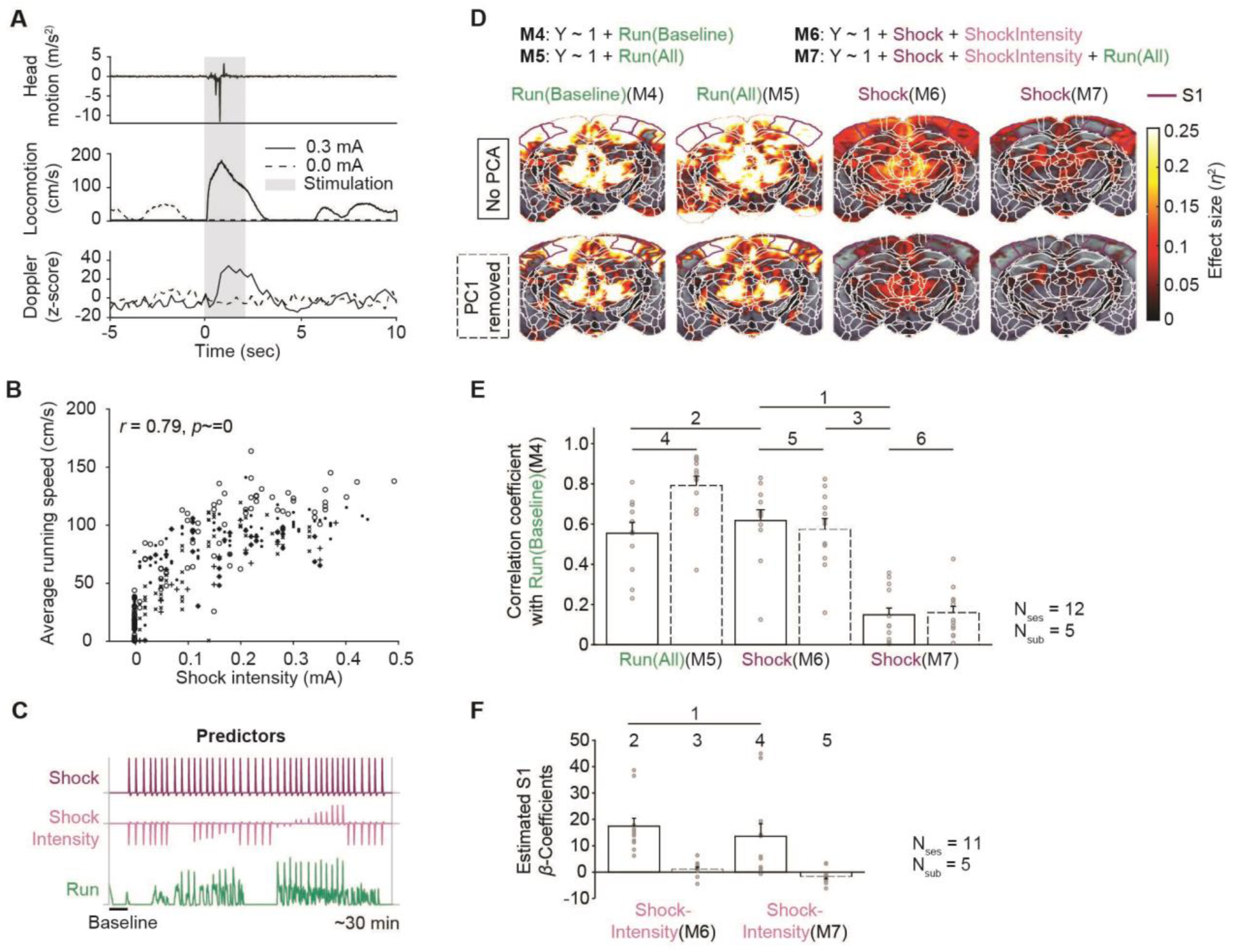
GLM modeling dissociates running from affective components of pain response. (A) Data traces of an example trial depicting the stimulation window (grey) of two shocks (0.0 mA, dashed lines; 0.3 mA, solid lines): Head-plate acceleration, running wheel velocity and the signal from one fUSI pixel from the primary somatosensory cortex (S1). (B) Average running speed during the shock period across shock intensities; dots represent trials, and shapes indicate animals. (C) Example predictors used in the GLM, representing all trials (Shock, purple), shock intensity (Normalised shockIntensity, pink) and Running (Run, dark green). Run refers to Running and Running⊗HRF combined. (D) GLM definitions using predictors from C, and example effect size maps of predictors across different GLMs without PC1 removal (top row) and with PC1 removal (bottom row). S1 is outlined in purple. (E) Group-level analyses showing 2D correlations of effect size maps between predictors from the different GLMs and correlations with the baseline running model without PC1 removal (solid lines) and with PC1 removal (dashed lines). (F) Estimated β-coefficients from S1 across the ShockIntensity predictor from the different GLMs without PC1 removal (solid lines) and with PC1 removal (dashed lines). Error bars represent standard deviation across sessions. Statistics represented in Table 2.

**Table 2.** Statistics of noxious stimulation experiment. The *t*-values are from two-sided paired t-tests corresponding to Figure 4EF. *BF_10_* indicates Bayes factors in favor of the alternative hypothesis. n = 12 for all comparisons in panel E. n = 11 for all comparisons in panel F.

| Comparison | Statistics |
| --- | --- |
| E1 | $t(11) = -6.23, p < 0.001, BF_{10} = 421$ |
| E2 | $t(11) = 1.21, p = 0.25, BF_{10} = 0.52$ |
| E3 | $t(11) = -5.65, p < 0.001, BF_{10} = 204$ |
| E4 | $t(11) = 5.44, p < 0.001, BF_{10} = 155$ |
| E5 | $t(11) = -1.76, p = 0.11, BF_{10} = 0.95$ |
| E6 | $t(11) = 0.21, p = 0.84, BF_{10} = 0.29$ |
| F1 | $t(10) = -1.58, p = 0.15, BF_{10} = 0.76$ |
| F2 | $t(10) = 5.47, p < 0.001, BF_{10} = 124$ |
| F3 | $t(10) = 1.39, p = 0.19, BF_{10} = 0.63$ |
| F4 | $t(10) = 2.74, p = 0.021, BF_{10} = 3.39$ |
| F5 | $t(10) = -1.96, p = 0.08, BF_{10} = 1.2$ |

In this experiment, due to the high correlation between running and noxious stimulation intensity, it was not possible to isolate responses to the noxious stimulation in the absence of movement and define a ground-truth map. Rather than using stationary trials as a reference for ground truth, we thus used a reference map that reflects spontaneous running-related activity, providing a reference estimate of running-related variance, as running serves as the primary source of noise we aim to eliminate. To create this reference map, we estimated running-related responses during the baseline period (2–7 min), before any shocks were administered (M4). By comparing models that include shock-related predictors to this baseline running model, the contribution of locomotion to the estimated responses can be quantified, allowing separation of shock-evoked neural activity from running-related activity.

Here, we qualitatively describe and illustrate how tail shock responses are captured by the different GLMs in a representative session (Fig. 4D). First, we examined whether running during the baseline period provides an appropriate proxy for running during shock administration by fitting a model of running across the entire session (M5). Second, we modeled all shock trials using a GLM that included a binary predictor indicating shock occurrence and a separate parametric predictor capturing shock intensity, enabling separation of shock-evoked responses from their scaling with stimulus intensity (M6). Consistent with earlier models (M2), the resulting responses reflected a mixture of stimulus-evoked and running-related activity, leading to elevated effect sizes across widespread regions (Fig. 4E, Shock, M6). Notably, the effect size map of the shock predictor in M6 overlapped substantially with the running predictors, especially in the somatosensory and mid-thalamic regions, indicating potential contamination by running-related variance in the variance explained by the shock stimulus. In contrast, the shock predictor in the final model (M7), which includes running as a covariate, showed spatial patterns distinct from the running predictors. Reduced effect sizes are observed in the retrosplenial cortex (RSC) and thalamic regions, whereas higher and more spatially localized effect sizes are observed in more localized areas such as primary somatosensory cortex (S1), which is associated with pain processing (Sun et al., 2023; Ziegler et al., 2023). We then proceeded to a quantitative analysis, employing similarity metrics to compare the models.

At the group level, we evaluated whether running during the baseline period provides an appropriate proxy for running during shock administration. The effect size maps of the running predictor across the entire session (M5) showed only a moderate similarity to the effect size maps derived from baseline running moderately (M4; *r* = 0.55 ± 0.18, Fig. 4E), suggesting that the running predictor from M5 contains variance beyond that observed during baseline running alone. This correlation increased after removing PC1 (*r* = 0.79 ± 0.16, Fig. 4E), suggesting the PC1 removal suppresses additional shared variance beyond motion-related effects in this case. When only modeling the shock activity without including running behavior (M6), the correlation with baseline running was similar for the Shock predictor from M6 and Running predictor from M5 (*r* = 0.62 ± 0.19. Fig. 4E), indicating substantial running-related contamination in the estimation of shock-related activity.

To understand how effectively motion-related activity during tail shock was separated in M7 at the group level compared to M6, we asked:

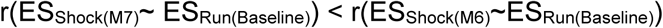

The Shock predictor estimated with M7 showed a markedly lower correlation with baseline running compared to M6 (*r* = 0.14 ± 0.14, Fig. 4E), demonstrating that M7 achieved significantly better separation of running-related variance.

To assess how PC1 removal affects the separation of different sources of variance in the data, the same analysis was performed after PC1 removing, yielding qualitatively similar effect size maps with enhanced spatial specificity across all models (Fig. 4D). Similar to the visual stimulation, removing PC1 does not improve the separation of running related responses from the shock response modeled in M6 (Fig. 4E), as evidenced by comparable correlations when comparing:

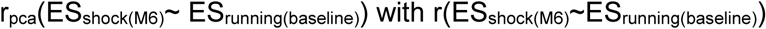

Incorporating running measurements in M7 continued to separate running-related variance in the data when PC1 removal was applied (*r* = 0.15 ± 0.12, Fig. 4E). Consistent with this, comparison of correlations between effect size maps of the Shock predictor in M7 with and without PC1 removal, relative to the baseline running reference, provided evidence in favour of the absence of a difference (Fig. 4E, BF<⅓) when comparing:

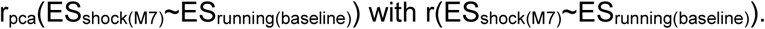

This indicates that explicitly modeling running renders the estimation of shock-evoked responses largely insensitive to PC1 removal (Fig. 4D). Finally, we directly compared separation achieved by GLM-based modeling versus PC1 removal by asking whether:

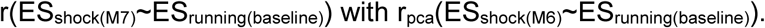

We found that the GLM approach achieves superior separation of running-related variance compared to PC1 removal alone (Fig. 4E).

To determine whether meaningful shock-evoked signals persist after accounting for running-related variance, we included Shock intensity as a predictor in the GLM (M6-7), allowing us to examine how variations in shock magnitude modulate activity conditional on running and other nuisance variables. We extracted the β estimate for the Shock intensity predictor from S1, a key region in pain processing (Sun et al., 2023; Ziegler et al., 2023). This β value reflects the strength of S1 activity specifically related to the magnitude of the shocks, providing a region-level measure of shock-evoked neural responses after accounting for motion and other nuisance variables (Fig. 4F). Relative to a model without running predictors (M6), adding running covariates (M7) yielded numerically smaller shock-intensity β values in S1, but the difference was not statistically significant (Fig. 4F). Importantly, both models produced positive and significant S1 modulation by the Shock intensity predictor (M6: *β* = 17.4; M7: *β* = 13.6, Fig. 4F), indicating that including behavior-informed covariates preserves interpretable shock-evoked responses. In contrast, removing PC1 during preprocessing markedly reduced S1 shock-intensity modulation (M6: *β* = 1.16, M7: *β* = -1.70, Fig. 4F), indicating that global component subtraction can remove variance that covaries with shock intensity and thus obscures stimulus-specific responses.

## Discussion

FUSI has demonstrated significant potential for mapping brain activity with high spatial resolution in rodents, providing an effective alternative to fMRI for measuring hemodynamic responses (Montaldo et al., 2022). While fUSI has been widely applied in controlled experimental paradigms, motion-related confounds remain the principal barrier to extending its use to more naturalistic paradigms. Our results show that explicit behavioral modeling, which includes continuous running speed predictors in the GLM, improves the fidelity of stimulus-evoked responses estimation in awake, headfixed mice. In a simple visual paradigm where ground-truth responses are well characterized and motion is minimal, we find that a model with behavioral covariates recovers response patterns that more closely match stationary-trial estimates than a model that omits them. In a noxious stimulation paradigm with strong locomotor and arousal components, the full model markedly reduces the correlation between shock-related and running-related response maps, indicating better separation of motion linked variance. Together, these findings support the utility of behavior-informed GLMs in both low- and high-motion paradigms.

Existing strategies to deal with motion-artifact mainly reduce motion contamination by exploiting image statistics, including SVD-based clutter filtering, frame/epoch censoring, and low-rank component removal (Brunner et al., 2021; Demene et al., 2015; Edelman et al., 2024; Urban et al., 2015). Recent benchmarking efforts, particularly for transcranial acquisitions where skull attenuation and macroscopic motion exacerbate artifacts, have begun to evaluate these approaches (Meur-Diebolt et al., n.d.-b). However, explicit behavior modeling is still absent. Our contribution complements these advances by making motion explicit in the statistical model: continuous behavioral measurements are entered as nuisance regressors in the GLM, aligning fUSI analysis with widely adopted fMRI practice for motion control. Rather than discarding data wholesale, we explain motion linked variance and retain interpretable task effects. The finding that stationary-trial responses are better approximated by the full model in the visual paradigm provides an intuitive validation (i.e., the model converges toward the low-motion “gold standard”). In the noxious stimulation paradigm, the strong correlation between baseline and session-wide running-effect maps indicates a stable, brain-wide footprint of locomotion, consistent with distributed hemodynamic and neural co-activation during movement. The shock map correlating of the reduced model with baseline running maps underscores how motion can masquerade as stimulus effects - a risk mitigated by the full model. Together, these results argue that behavior-informed GLMs are a useful last mile after standard SVD filtering/clutter rejection and selective censoring.

Mechanistically, motion affects fUSI signals via at least three intertwined pathways: (i) mechanical displacement that globally perturbs the acoustic field and backscattered power; (ii) systemic physiology (elevated cardiac output/arterial pulsatility during locomotion) that modulates cerebral blood volume; and (iii) movement-related neural activation that co-varies with behavior. Treating motion as measured covariates allows the model to absorb variance from these sources, rather than relying solely on image-intrinsic signatures. This complements SVD filters - which exploit spatiotemporal coherence differences between tissue and flow (Demene et al., 2015) but cannot know when the animal ran - and avoids the data loss inherent to aggressive censoring (Brunner et al., 2021).

Importantly, our conclusions remain largely consistent even when the first principal component is removed, indicating that behavioral predictors provide information not redundant with global component suppression. This is particularly relevant because PC1 removal is used in several awake fUSI pipelines as a pragmatic step when unspecified movement artifacts are suspected (Brunner et al., 2021; Edelman et al., 2024; Siegenthaler et al., 2025). In specific experimental conditions like noxious stimulation that involves intense motion, we showed that the GLM approach managed to disassociate the response pattern of movement and maintain a meaningful level of stimulus evoked response in the key region of interest while simply removing PC1 does not, suggesting the behavior modeling approach overperform the PCA approach in some circumstances. By validating across low-motion (visual stimulation) and high-motion (noxious stimulation) regimes, we show that behavioral covariates remain beneficial across different signal-to-confound ratios - a key criterion for naturalistic paradigms.

fUSI’s portability, mesoscopic resolution coverage (100 µm / 100 ms) and brain-wide coverage enable mapping in awake animals and compatibility with electrophysiology and optogenetics (Brunner et al., 2021; Edelman et al., 2021; Macé et al., 2011; Nunez-Elizalde et al., 2022). By decreasing motion-linked contamination without discarding data, behavior-informed GLMs make it more practical to study self-generated behaviors (running, orienting), state changes (arousal) and more complex situations involved with virtual reality or social interaction in head-fixed preparations. This approach bridges the gap between rigid, stimulus-locked tasks and richer behavioral contexts, and it aligns with ongoing efforts to stabilize chronic access and standardize artifact handling (Edelman et al., 2024; Meur-Diebolt et al., n.d.-b).

Our study has limitations. First, as in fMRI, motion covariates can correlate with task timing, risking over- or under-correction of true effects. This can be mitigated by (i) including derivatives/lags of running/head motion, (ii) orthogonalizing predictors when appropriate, and (iii) reporting variance-partitioning (variance explained by motion vs. task). Second, systemic physiological changes (e.g., respiration, heart rate) are only partially captured by running speed/head motion, adding physiology channels could further improve separation. Third, GLMs assume linearity/additivity, future work could test nonlinear terms or state-dependent models. Finally, while we validated against stationary-trial responses and across two paradigms in 2D imaging, broader cross-task and cross-region generalization in 3D imaging (Bertolo et al., 2021; Gesnik et al., 2017) and replication across laboratories will strengthen confidence in our approach.

Our findings indicate that in awake, head-fixed rodent experiments, incorporating continuous behavioral covariates as GLM nuisance regressors can improve the precision of stimulus-response estimates in low-motion contexts and materially help reduce motion confounds in high-motion contexts - without disproportionate data loss. In combination with standard clutter filtering and pragmatic censoring, behavior-informed GLMs provide a practical path toward more naturalistic fUSI in head-fixed rodents and align fUSI analysis with well-established practices in the fMRI community. In freely moving animals, overall motion may increase, but because the probe moves with the subject, we do not expect the approach to differ substantially, with performance mainly depending on how movement correlates with the state of interest. As the field moves toward richer behaviors and transcranial or freely moving preparations, explicit modeling of behavior should become part of the default fUSI data analysis.

## Materials and Methods

### Contact for Resource Sharing

Further information and requests for resources should be directed to and be fulfilled by the Lead Contact, Valeria Gazzola.

### Animals

5 female C57BL/6JRj mice (7-12 weeks old) were obtained from Janvier, France. Upon arrival, the animals were socially housed at room temperature (22-24°C), on a reversed 12:12 light:dark cycle (lights off at 07:00). Food and water were provided ad libitum. All experimental procedures were pre-approved by the Centrale Commissie Dierproeven of the Netherlands (AVD8010020209725) and the animal welfare body of the Netherlands institute for Neuroscience (IVD, study dossier 233602).

### Handling

The mice were handled for 5-15 minutes per day, minimally for three days, one or two times per day, until they were comfortable with the experimenters.

### Cranial window surgery

Mice (13-15 weeks; 20-23 grams) were anesthetized with 4-5% isoflurane (induction) and ∼2% isoflurane (maintenance) in oxygen (0.8L/min flow rate) through a nose cone. After induction, mice received carprofen (5 mg/kg, s.c.) and buprenorphine (0.1 mg/kg, s.c.) for analgesia, and dexamethasone (8 mg/kg, s.c.) to prevent cerebral inflammation. Analgesics were administered at least 20 minutes prior to the first incision to ensure adequate onset of action. Following the initial incision, lidocaine was applied to the periosteum to provide local anesthesia. A craniotomy (maximally -4.5 to 0 AP, -5 to +5 ML, in order to include the somatosensory cortex (S1), lateral geniculate nucleus (LGN) and lateral posterior nucleus of the thalamus (LPN)) was drilled, while the dura was kept intact. After removal of the skull, the dura was covered with artificial dura. A flap of TPX (Polymethylpentene, 250 um thick) was cut to fit the craniotomy, positioned on top of the artificial dura and the bone and sealed. A custom headplate was attached to the skull with tissue adhesive and dental cement. To keep the weight on the animal’s head below 10% of its body weight, the headplate was made of titanium, weighing 1.2 grams. After cleaning the wound edges with saline and Hibicet, Prontosan was applied to control the bacterial burden. After the surgery, mice were placed in a heated recovery chamber and given additional buprenorphine (0.1 mg/kg s.c.) 4-6 hours after the first injection. After full recovery from anesthesia, mice were transferred from the recovery chamber to the home cage. Post-operative analgesia included daily Rimadyl injections (5 mg/kg) and diet gels until 48-72 hours post-OP (depending on the recovery). Mice recovered for one full week before being trained on the set-up.

### Habituation to head fixation

To reduce any stress possibly associated with the head-fixation procedure or the set-up, mice were extensively habituated to the head-fixed experimental context (Figure 1A). Mice were trained maximally twice a day for 6-12 days, while increasing the time of the fixation over the sessions (from min 10 to max 90 min per session per day). The running wheel was cleaned with lemon dishwashing soap and the probe was cleaned with disinfectant wipes. Then, mice were head-fixed on the running wheel and the acoustic window was cleaned with 70% ethanol. Acoustic gel was applied to allow for ultrasound coupling and the ultrasound probe was positioned into the gel 2 mm above the cranial window implant. Mice were habituated to the electrodes on their tail by placing them 5 mm proximal and distal of the tail (Dolensek et al., 2020). To ensure proper electrical conduction, electrode gel was applied between each electrode and the tail, while preventing direct contact between the two electrodes through the gel. Moreover, mice were habituated to the visual stimuli projected on the monitor ∼20 cm in front of them. Dim-light conditions were applied. After habituation, mice were handled for five minutes and returned to their home cage. The running wheel and probe were cleaned after each animal to prevent contamination and to remove stress pheromones.

### Visual stimulation

The medium for presentation of visual stimuli was a 24 inch monitor (U2410, Dell, USA), placed ∼20 cm in front of the animal, facing it perpendicularly (Fig. 1A). During the baseline, a black screen was displayed for 2-5min. Then, 5-10 stimuli (black and white moving grating randomly shown in eight angles, switching at 2Hz) were on for 15 seconds with a 45 seconds inter-stimulus interval (ISI) showing a black screen (Fig. 1C).

### Noxious stimulation

Noxious stimuli were delivered using a programmable shock generator (ENV-414S, Med Associates Inc., VT, USA) equipped with output monitoring and a dual-electrode setup to ensure stable tail contact. Mice received three shocks per intensity level, with current amplitudes increasing in 0.05 mA increments from 0.05 mA to 0.4 mA. Blocks of five control trials (0.0 mA) were delivered before the first shock, midway through the session (after the 0.2 mA shocks), and after the final shock (Fig. 1C). The inter-trial interval (ITI) between the shocks was 40 ± 10 seconds.

### Recording motion

Two external sensors were added that quantify motions. Firstly, a miniature ±2g MEMS accelerometer with 0.24mg/digit resolution is rigidly mounted on the head-plate holder to flag frames contaminated by true head motion. Secondly, a custom, non-slip treadmill (0.05 cm sensitivity) records instantaneous running speed while allowing voluntary locomotion, which reduces stress and provides a continuous behavioural read-out.

### fUSI data acquisition

To ensure consistent positioning of the imaging plane across sessions (Fig. 1B), the fUSI probe was mounted on a precision motorized stage allowing rotation and fine-scale translation in all spatial directions. The head plate could be tilted by ±10° in both anteroposterior and mediolateral axes to compensate for minor variations in surgical placement of the head-plate. The imaging plane was selected through an iterative three-step procedure: (1) acquisition of a volumetric scan across the craniotomy, (2) registration of the scan to the Allen Brain Atlas and (3) selection of the corresponding slice within the atlas framework. These steps were performed using established MATLAB tools (Brunner et al., 2021) and custom analysis scripts. Once optimal alignment was achieved, probe position and orientation were saved for subsequent sessions to ensure reproducible imaging across experimental sessions within one animal.

To evaluate the capabilities of our system, we aimed to image relatively small, deep brain regions in both experiments. For the visual stimuli, we selected the LGN and LPN, which have previously been shown to respond to visual stimuli using fUSI (Brunner et al., 2020). S1 was targeted in the noxious stimulation experiment for its role in processing nociceptive signals and its involvement in pain perception (Sun et al., 2023; Ziegler et al., 2023).

The fUSI data was acquired using an ultrasound transducer (L22-14vX; center frequency 18MHz, bandwidth 60%) connected to a high–frame rate ultrasound scanner (Vantage, Verasonics, USA). The system transmitted 12 angled plane waves at 12 kHz pulse repetition frequency, which were compounded to B-mode images, resulting in a B-mode frame rate of 1 kHz. Doppler ensembles of 200 frames (200 ms) were used to capture hemodynamic activity. Static tissue signals were removed using a singular value decomposition (SVD) clutter filter with a fixed threshold of removing the first 50 singular vectors (Demene et al., 2015). Power Doppler Images (PDIs) were subsequently formed from the clutter filtered data. Real-time reconstruction and filtering were performed on the acquisition workstation using CUDA-based processing, allowing continuous Power Doppler imaging at 5 Hz.

### Multimodal data acquisition and alignment

To integrate neural, behavioral, and stimulation data, all experimental signals were synchronized and temporally aligned using a TTL-based framework. The Verasonics scanner was connected to a dedicated computer for fUSI data acquisition, while a separate computer controlled experimental protocols and logged auxiliary signals (Fig. 5). The experimental computer controlled recording and stimulation devices through a digital input–output (DIO) interface by sending transistor–transistor logic (TTL) signals to the corresponding hardware, while recording timing information by sending additional TTLs to the data acquisition (DAQ) system. Recording devices measured behavioral outputs (accelerometer, running wheel), whereas stimulation devices delivered stimuli to the animal (monitor, shock electrodes). Data streams from all recording devices and the timestamps of all TTL events recorded by the DAQ were saved on the experimental computer for subsequent alignment. To synchronize the power Doppler images to the other behavioral readouts, the trigger out of the Verasonics was connected to the DAQ and configured to send a trigger at the start of a Doppler ensemble. Since the Verasonics TTL trigger is a 1 us pulse, but the DAQ sampled at 5kHz, the TTL triggers were extended to 5 ms using an Arduino Uno R3 to avoid missing triggers. The experimental PC also controlled the linear motor (Zaber, X-DMQ12P), enabling both automated and manual positioning of the ultrasound probe.

**Figure 5.**
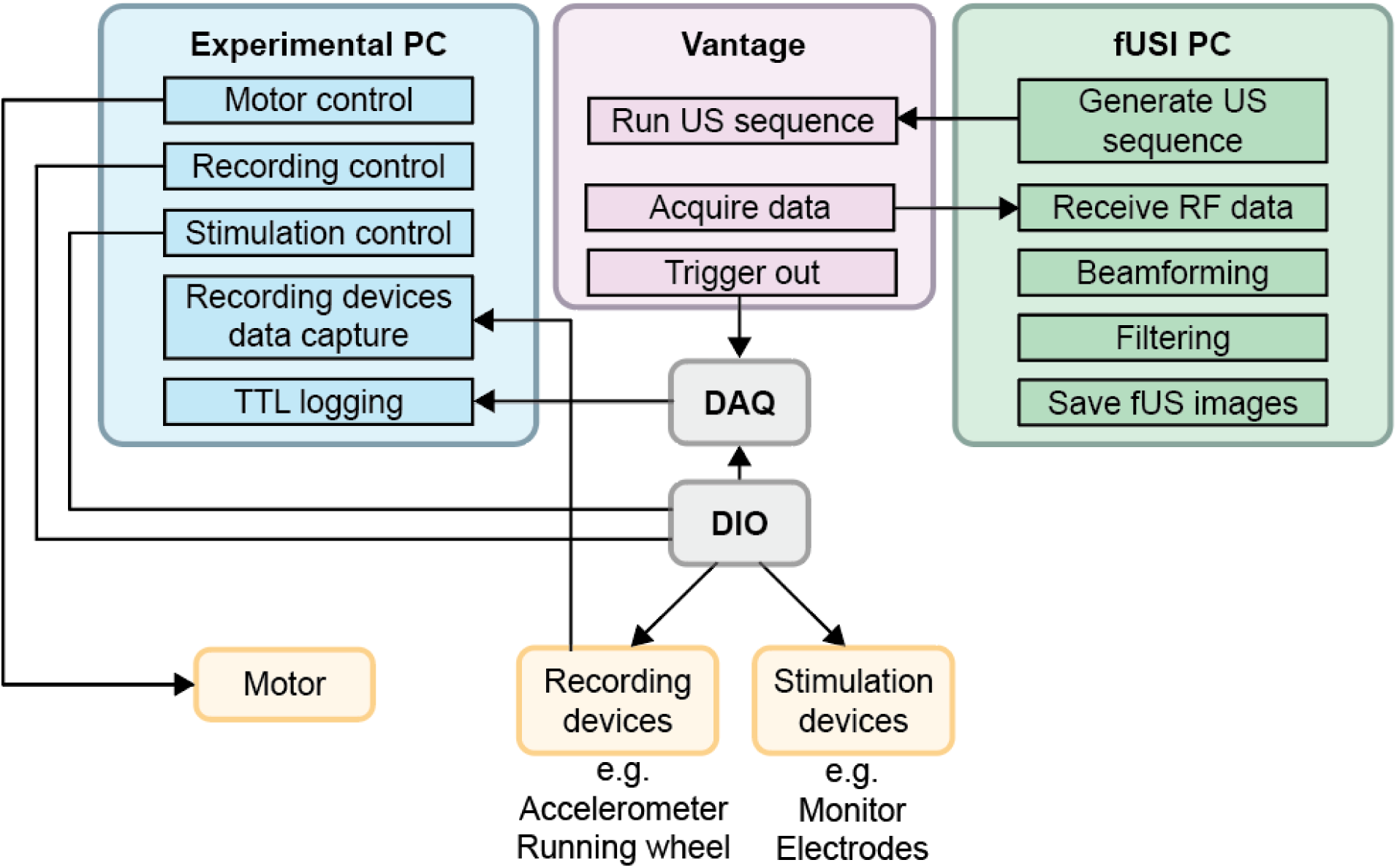
Experimental set-up. The system consists of several main hardware components. The experimental PC (blue) serves as the central control unit, managing the positioning of the motor and the control of the recording and stimulation devices (yellow) while capturing the TTLs and the data from the recording devices. The alignment is maintained via the DIO and DAQ (grey), which transmit digital signals from the experimental computer to the recording and stimulation devices, and relay timing data back to the PC. The fUSI system consists of (1) the fUSI PC (green), which generates imaging sequences, acquires raw data, and processes it into fUSI images and (2) the Vantage system (purple), which controls the fUSI probe and provides a trigger for each frame.

### Preprocessing of fUSI data

First, we resampled the fUSI signal to a uniform 5 Hz sampling rate and calculated the percentage change of the signal using the mean value of the whole recording in each pixel. Next, we applied a discrete cosine transformation (DCT) highpass filter with a cutoff frequency of 0.002 Hz to remove very slow baseline drifts in the fUSI signal (e.g., probe coupling, temperature- or physiology-related trends) while preserving task-related and behavioral fluctuations. Head motion was quantified as the vector magnitude of the 3-axis acceleration recorded on the head plate: 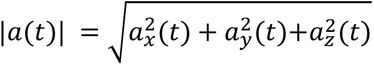 and resampled to the same sample rate as the fUSI signal for artifact detection. A frame-level artifact detection procedure was performed on a per-pixel, per-frame basis using two complementary criteria: First, a frame-wise criterion flagged complete frames in which the head motion magnitude exceeded the session-wise threshold (z(|*a*(*t*)|) > 3.5). Second, a PDI-based criterion flagged, for each pixel separately, frames in which the fUSI signal exceeded a pixel-specific threshold (z(fUSI(t)) > 5). A frame was labeled as an artifact for a given pixel if it met either criterion. Labeled samples were removed independently for each pixel and replaced by linear interpolation across time. Then the fUSI signal was spatially smoothed with a Gaussian filter of 1 sigma (FWHM ≈ 235.5 μm horizontal, 117.8 μm vertical).

To quantify temporal signal stability in the fUSI time series, we computed pixel wise temporal signal-to-noise ratio (tSNR). For each pixel, classical tSNR was calculated as the mean signal over time divided by its standard deviation.

### Removal of locomotion-related confounds

When animals walk or run, this can create two sources of signal that are not directly task related. First, physical motion of the brain will affect the Doppler signal instantaneously. Second, brain activity associated with running will trigger hemodynamic activity with the delay of the hemodynamic response function. To account for both components in each pixel of the fUSI signal, we use a General Linear Model approach including different transformations of the measured running speed (M_Running_). The raw running speed (Running) and its first derivative (acceleration, Running’) were included to model the physical motion, whereas the raw speed convolved with the hemodynamic response function (Nunez-Elizalde et al., 2022), *Running* ⊗ *HRF,* was included to model brain activity associated with running. We used a canonical fUSI hemodynamic response function as characterized previously (Nunez-Elizalde et al., 2022), which provides a reasonable approximation of the temporal dynamics of blood-volume changes measured with fUSI. The residual (ε) represents the portion of the signal not related to motion.

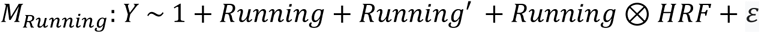

These variables are also incorporated in GLMs in the following sections to account for the contribution of running in the fUSI signal when modeling the experimental tasks. The *Running*′ predictor is omitted in subsequent GLMs due to its low percent of variance explained in the data.

To characterize dominant spatiotemporal patterns in the fUSI signal, we performed a pixel-wise principal component analysis (PCA). The 3-D data (height × width × time) were reshaped into a two-dimensional matrix of size T × nPixel, where T number of time points, nPixel is the number of pixels. Then, the time series of each pixel was z-scored, so that the subsequent correlation-based PCA would capture shared temporal patterns independent of individual pixel amplitude. We then applied economy-size SVD to the resulting matrix to extract dominant spatial patterns and their associated time courses. Spatial maps of the first principal component were obtained by reshaping the pixel loadings to the original 2-D grid.

To evaluate the effect of removing PC1 on motion-related artifact correction, we performed all analyses both with and without PC1 from the high-pass–filtered fUSI signals. Removal of PC1 was implemented by projecting its spatial map and associated time course out of the data matrix, following the procedure described in (Edelman et al., 2024). Only the first principal component was removed, as it captured the dominant shared variance across pixels; higher-order components were not removed to avoid eliminating spatially specific neural signals.

### Estimating neural activity during visual stimulation using GLM and Pearson correlation

To validate the effectiveness of the GLM approach in controlling the influence of running during visual stimulation, we defined three GLMs (Fig. 3C): M1 characterizes visual stimulation-induced activity during non-running trials and serves as a reference model. M2 models activity related to all visual stimulations without taking running into account. M3 represents activity related to all visual stimulations while explicitly modeling running speed and its interaction with visual stimulation as separate regressors, thereby isolating visual-stimulus–related activity. In all models, the visual stimulation predictor (Visual) was constructed as a boxcar function with value 1 while stimulus is on and 0 while off, convolved with a hemodynamic response function (Nunez-Elizalde et al., 2022). Predictors convolved with the HRF are denoted with the ⊗HRF notation.

M1 served as a reference response pattern to visual stimulation. Only trials that were not confounded by running were included, defined as trials in which wheel velocity exceeded 2 cm/s for less than 200ms during the stimulation period. In some early sessions, where animals moved more, no trials met this criterion; these sessions were excluded from further analysis(n=7). This resulted in a total of 27 sessions from 5 animals (see Table S1).

The model was defined as:

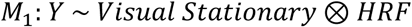

where Y represents the pixel-wise fUSI response.The intercept and residual term are omitted here and for all subsequent models for clarity.

Next, M2 captures activity related to all visual stimuli without accounting for running-related variability, which may confound the estimation of visual-evoked brain activity. The model was defined as:

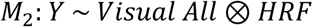

Lastly, M3 estimates visual responses to all stimuli, as in M2, but is designed to dissociate visual-evoked activity from running-related activity. The model includes running speed as a predictor in two forms. The un-convolved predictor (Running) accounts for immediate movement-related signals, while the HRF-convolved predictor (*Running* ⊗ *HRF*) captures running-related neural activity. To capture neural processing of visual stimuli that is modulated by running, an interaction term between Visual and Running was included (Troncoso et al., 2015). This interaction was computed by multiplying the predictors before convolution with the HRF. The model was defined as:

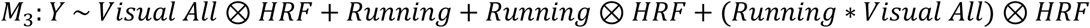

To quantify the contribution of each predictor to pixel-wise fUSI responses, the GLMs were fitted using ordinary least squares regression for each pixel in each individual session. For each predictor, the partial eta squared (η^2^_p_) was calculated to quantify the effect size.

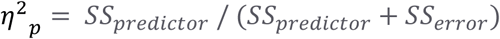

As an alternative approach, we calculated Pearson correlations between the observed responses Y and the predictors (Visual Stationary⊗HRF and Visual All⊗HRF) to quantify how strongly each pixel’s activity tracks the stimulus.

To assess the spatial similarity between GLM and correlation, 2D correlations were calculated between the effect size maps and the correlation maps. To statistically evaluate differences between the maps across sessions, a paired sample t test was applied across all 27 sessions (from 5 animals). Where appropriate, Bayes factors of equivalent test were additionally computed to quantify evidence for an effect (BF >3) or for the absence of an effect (BF < ⅓; Keysers et al., 2020).

### Contrast to noise ratio

To evaluate the detectability of stimulus-evoked signals in our imaging system, we computed the contrast-to-noise ratio (CNR) for each stimulation epoch within each region of interest (ROI). The averaged time series were aligned to stimulus onset and two windows were defined: a baseline window (−5 to 0 s before onset) and a response window (0 to +15 s after onset). CNR was calculated using a variance-based metric:

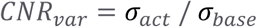

where σ_act_ is the standard deviation of the fUSI signal within the response window and σ_base_ is the standard deviation in the baseline window. CNR was computed for each trial and then averaged across trials within a session. Differences in CNR across analyses methods were assessed using paired sample t-tests across all sessions (N depends on the number of sessions with coverage of the specific ROI).

### Estimating neural activity during noxious stimulation using GLM

To evaluate the performance of the GLM approach, we applied it to separate shock-evoked neural activity from running-related signals using four GLMs (Fig. 4C): M4 characterizes running related signals during the baseline of the session and serves as a reference model. M5 captures running-related activity throughout the entire session. M6 represents activity related to all noxious stimulations without taking running into account. M7 depicts activity related to all noxious stimulations while explicitly modeling running, thereby isolating noxious-stimulus–related activity.

M4 gives a reference map of running activity during the baseline before any shock administration. The model was fitted as:

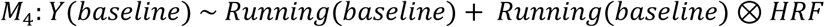

M5 is similar to M4, but captures running-related activity throughout the session, including during shocks:

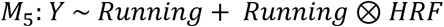

M6 captures shock-evoked responses without accounting for running. Shock events were encoded as the same boxcar function with value 1 while stimulus is on and 0 while off, regardless of stimulus intensity. This time series was then convolved with the HRF to model the expected fUSI signal (*Shock HRF*). A complementary predictor, the shock intensity predictor, was constructed by replacing all 1s with the z-scored actual stimulus intensity delivered to the animal, and then it was similarly convolved with the HRF (*Shock Intensity* ⊗ *HRF*).

The model was fitted as:

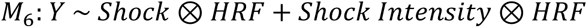

M7 aims at dissociation of shock-related brain activity from running-related activity. The model was defined as:

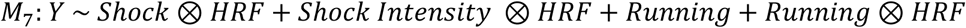

The GLMs were fitted using ordinary least squares regression in the same manner as for the visual experiment. For each pixel, the partial eta squared (η^2^_p_) was calculated to quantify the effect size of each predictor. Differences between correlation coefficients of the effect size maps across models was assessed using paired sample t-tests across all 12 sessions (from 5 animals).

To quantify how specific regions correlate with the shock intensity predictor, β-coefficients were extracted per ROI by averaging the β values of all pixels within each ROI for a specific GLM. Differences between β values across models were tested using paired sample t-tests across all sessions (N depends on the number of sessions with coverage of the specific ROI).

## Author contributions

C. Qin, F. Nelissen, B. Heiles, C. Keysers and V. Gazzola contributed to conceptualization. Methodology was developed by F. Nelissen, R. Waasdorp, B. Heiles, C. Rojas, L. deAngelis, C. Qin, M. Heemsterk, C. Keysers and V. Gazzola. Software was provided by R. Waasdorp, C. Qin, C. Rojas, and L. deAngelis, with validation by F. Nelissen, R. Waasdorp, C. Rojas, and L. deAngelis. Formal analysis was performed by C. Qin, F. Nelissen, C. Keysers and V. Gazzola, and investigation was carried out by F. Nelissen and A. Lotfi. Resources were provided by V. Gazzola, P. Kruizinga, and D. Maresca, with data curation by F. Nelissen, C. Qin and R. Waasdorp. The original draft was written by C. Qin, F. Nelissen, and B. Heiles, with review and editing by all authors. Visualization was performed by F. Nelissen and C. Qin. Supervision was provided by F. Nelissen, B. Heiles, C. Qin, C. Rojas, L. deAngelis, V. Gazzola and D. Maresca. Project administration was led by F. Nelissen, and funding acquisition was secured by V. Gazzola, B. Heiles and D. Maresca.

## Acknowledgments

The authors thank B. Koekkoek, B. Generowicz and P. Kruizinga for providing the initial version of the fUSI acquisition system. Further, the authors thank A. de Groot and M. Vink for building the accelerometer and tail shock device, J. Veldhuijzen and S. Super for building the running wheel and head-post, and J. Wekselblatt for guidance on surgical procedures and the head-plate design. The authors also thank I. da Silva Serra for guidance on surgical procedures, R. van Tol for supporting the 3D setup visualization, S. Voges for contributing to the piloting of the tail shock device, A. Babiczky for advice on imaging planes, and R. Rajendran for writing ethical pre-approval AVD8010020209725. Finally, the authors thank V. Galligioni, G. de Fluiter, and veterinary staff of the Netherlands Institute of Neuroscience for their continuous support. ChatGPT was used as a writing assistant in preparing this manuscript.

