## Supplementary material for "Modeling behavior to disentangle motion-related effects in functional ultrasound imaging in awake, head-fixed mice"

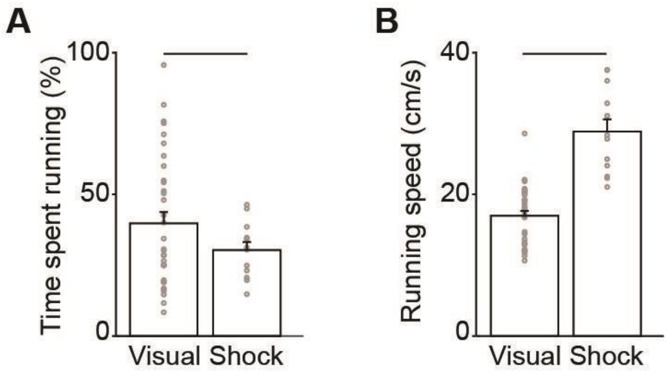


**Supplementary Figure S1. Running behavior quantification.** (A) Percentage of time spent running in visual and shock stimulation sessions, including baseline. (B) Average running speed during visual and shock stimulation sessions, including baseline. Error bars represent standard deviation across sessions. Statistics represented in Supplementary Table 2.

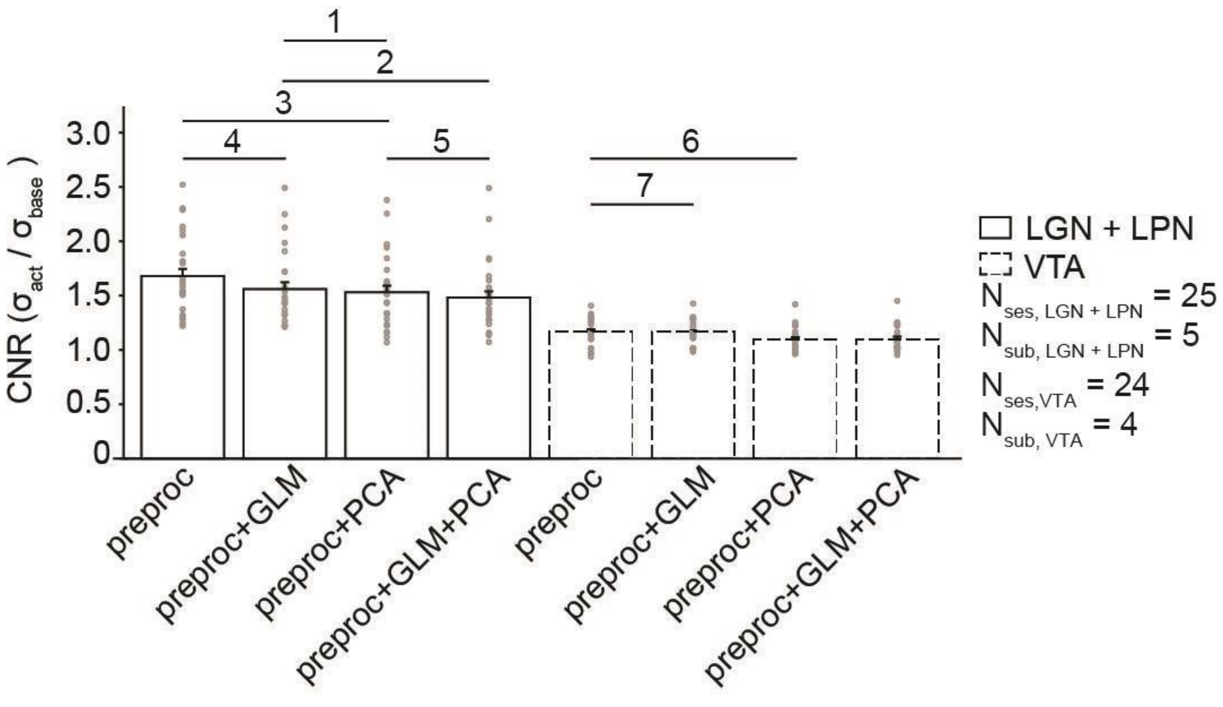


**Supplementary Figure S2. Effect of motion correction on Contrast to Noise Ratio (CNR).** Variance based CNR in visual regions (LGN+LPN, solid lines) and a control region (VTA, dashed lines) is shown for GLM, PCA, and combined corrections after preprocessing steps. Error bars represent standard deviation across sessions. Statistics represented in Supplementary Table 3.

**Supplementary Table 1. Information of all included experimental sessions**

| Session type | Animal Index | Number of trials | Number of running trials | Baseline duration (s) |
|---|---|---|---|---|
| Visual | 2 | 10 | 10 | 615 |
| Visual | 2 | 10 | 4 | 131 |
| Visual | 3 | 10 | 10 | 132 |
| Visual | 3 | 10 | 10 | 135 |
| Visual | 4 | 10 | 7 | 137 |
| Visual | 4 | 10 | 6 | 135 |
| Visual | 5 | 10 | 10 | 134 |
| Visual | 5 | 10 | 10 | 611 |
| Visual | 1 | 5 | 3 | 612 |
| Visual | 2 | 5 | 5 | 131 |
| Visual | 1 | 10 | 9 | 135 |
| Visual | 3 | 5 | 4 | 129 |
| Visual | 4 | 5 | 3 | 311 |

| | | | | |
|---|---|---|---|---|
| Visual | 5 | 5 | 2 | 314 |
| Visual | 1 | 10 | 7 | 309 |
| Visual | 2 | 10 | 6 | 608 |
| Visual | 2 | 10 | 6 | 618 |
| Visual | 3 | 10 | 10 | 611 |
| Visual | 3 | 10 | 9 | 615 |
| Visual | 3 | 10 | 8 | 610 |
| Visual | 3 | 10 | 9 | 611 |
| Visual | 3 | 10 | 4 | 617 |
| Visual | 3 | 10 | 9 | 613 |
| Visual | 3 | 10 | 4 | 613 |
| Visual | 4 | 10 | 5 | 610 |
| Visual | 4 | 10 | 5 | 618 |
| Visual | 4 | 10 | 5 | 613 |
| Visual | 4 | 10 | 5 | 616 |

| | | | | |
|---|---|---|---|---|
| Visual | 4 | 10 | 7 | 613 |
| Visual | 4 | 10 | 5 | 611 |
| Visual | 5 | 20 | 11 | 612 |
| Visual | 5 | 20 | 12 | 630 |
| Visual | 2 | 30 | 9 | 613 |
| Visual | 5 | 30 | 27 | 615 |
| Shock | 2 | 40 | 20 | 132 |
| Shock | 1 | 40 | 28 | 132 |
| Shock | 1 | 40 | 32 | 132 |
| Shock | 1 | 42 | 33 | 431 |
| Shock | 3 | 42 | 30 | 430 |
| Shock | 3 | 42 | 27 | 436 |
| Shock | 3 | 42 | 36 | 436 |
| Shock | 4 | 42 | 30 | 429 |
| Shock | 4 | 42 | 29 | 431 |

| | | | | |
|---|---|---|---|---|
| Shock | 5 | 42 | 26 | 435 |
| Shock | 5 | 42 | 26 | 453 |
| Shock | 5 | 42 | 30 | 445 |

**Supplementary Table 2. Statistics of running behavior quantification**

| Comparison | Statistic |
|---|---|
| A | $t(44) = -1.36$, $p = 0.18$, $BF_{10} = 0.66$ |
| B | $t(44) = 7.57$, $p < 0.001$, $BF_{10} = 4.11 * 10^6$ |

**Supplementary Table 2.** The *t*-values are from two-sided paired t-tests corresponding to Figure S1AB. $BF_{10}$ indicates Bayes factors in favor of the alternative hypothesis. n = 45 for all comparisons.

**Supplementary Table 3. Statistics of effects of motion correction on Contrast to Noise Ratio (CNR)**

| Comparison | Statistic |
|---|---|
| 1 | $t(24) = -0.94$, $p = 0.13$, $BF_{10} = 0.28$ |
| 2 | $t(24) = -3.09$, $p < 0.001$, $BF_{10} = 8.19$ |
| 3 | $t(24) = -3.90$, $p < 0.001$, $BF_{10} = 48.2$ |
| 4 | $t(24) = -5.05$, $p < 0.001$, $BF_{10} = 677$ |
| 5 | $t(24) = -2.88$, $p < 0.001$, $BF_{10} = 5.35$ |
| 6 | $t(20) = -4.34$, $p < 0.001$, $BF_{10} = 102$ |
| 7 | $t(20) = -0.56$, $p < 0.001$, $BF_{10} = 0.21$ |

**Supplementary Table 3.** The *t*-values are from two-sided paired t-tests corresponding to Figure S2. $BF_{10}$ indicates Bayes factors in favor of the alternative hypothesis. n = 25 for comparisons 1-5 and n = 21 for comparisons 6 and 7.
